# Uncertainty-aware quantitative analysis of high-throughput live cell migration data

**DOI:** 10.1101/2025.06.12.659342

**Authors:** Simo Kitanovski, Shannon Conroy, Justin Sonneck, Madeleine Dorsch, Sebastian Urban, Jianxu Chen, Barbara M. Grüner, Daniel Hoffmann

## Abstract

Accurate quantification of cell migration velocity is essential for understanding biological processes such as development, immune function, and cancer metastasis. High-throughput migration assays generate complex, hierarchically structured datasets with technical noise, batch effects, and biological variability, introducing uncertainty into velocity estimates that current methods often fail to quantify. To address this, we present cellmig, a computational tool using Bayesian hierarchical modeling to separate biological signals from technical variation while explicitly quantifying uncertainty in migration velocity. cellmig provides a robust framework for analyzing migration assays, including dose-response studies and large-scale screens with biological and technical replicates. By modeling biological variability and technical confounders within a unified Bayesian framework, it estimates condition-specific effects (e.g., drug effects) on cell velocity with probabilistic uncertainty intervals, avoiding common pitfalls of null-hypothesis testing. Its generative models simulate migration under various assumptions, aiding experimental planning. Overall, cellmig improves reproducibility and comparability across studies, offering deeper biological insight.

## Introduction

Cell migration is essential for many vital biological functions, for instance division or differentiation, and it impacts pathological processes including cancer metastasis (Aman and Piotrowski, 2010; Friedl and Gilmour, 2009; Luster et al., 2005; Reig et al., 2014). Diverse experimental methods have been developed aimed at quantifying different facets of cellular migration, including velocity, path linearity, average turning angle, time persistence or directional bias. These methods depend on a variety of variables, such as the cell type, exposure to chemical or physical stimuli and cell-cell interactions (Toscano et al., 2024).

The most common method for studying cell migration involves tracking cells with time-lapse microscopy followed by the acquisition of images at fixed time intervals over several hours (Masuzzo et al., 2016; Toscano et al., 2024). A high-throughput version of this method facilitates the analysis of cell velocity across dozens of wells treated under different biological conditions (Cibir et al., 2023; Kramer et al., 2013). This is followed by computational image processing to extract quantitative cell velocities and to detect changes following perturbations such as drug treatments (Cibir et al., 2023; Hulkower and Herber, 2011). However, these workflows generate considerable technical noise (e.g., imaging artifacts, unprecise pipetting) and biological variability (e.g., cell-to-cell heterogeneity), and therefore technical and biological replicates are necessary, leading to large, hierarchically structured datasets: cells are nested within technical replicates that are nested within biological replicates (Lee et al., 2021; Masuzzo et al., 2013).

Current statistical analyses of such data rely on qualitative approaches, see e.g. Figure 3B from Masuzzo et al. (2016), or inadequate statistical methods, e.g., the nonparametric Wilcoxon or Kruskal-Wallis rank-sum tests (Kleino et al., 2025; Masuzzo et al., 2013; Shannon et al., 2024; Wiggins et al., 2023; Wortel et al., 2021). These approaches usually ignore the hierarchical structure of the data and fail to explicitly quantify uncertainty arising from technical or biological variability, reducing reproducibility. Similarly, to graphically represent the data, biologists typically quantify the velocity of cells from a single experiment, depict them in simple plots for each individual cell of that particular experiment and then describe the additional biological replicates without providing a graphical representation of the entire dataset (Dorsch et al., 2021). To address this gap, we present cellmig, an R package implementing Bayesian hierarchical models for migration analysis. cellmig quantifies condition-specific velocity changes (e.g., drug effects) while modeling nested data structures and technical artifacts, providing uncertainty-aware estimates through credible intervals. It further enables synthetic data generation for experimental design optimization. By integrating these features into an accessible tool, cellmig enables quantitative research and enhances reproducibility. We demonstrate this by analyzing our own cell migration assays with over 30,000 cells and 20 biological conditions.

## Materials and methods

### Cell line and cultures

ASPC1 cells (ATCC Cat# CRL-1682, RRID:CVCL_0152, sex: female) were cultured under humified atmosphere at 37 ℃ and 5% CO_2_ in DMEM high glucose, pyruvate (Gibco, Cat # 41966052) supplemented with 10% FBS (VWR/Biowest, Cat #S1600-500), 1% non-essential amino acids (ThermoFisher, Cat #11140035), 50 U/ml penicillin and 50 µg/ml streptomycin (ThermoFisher, Cat #10378016). Identity was authenticated by STR analysis. Testing to exclude mycoplasma contamination was performed regularly.

### Chemicals

The following commercially available compounds were used at concentrations as indicated: fluvastatin (abcam, Cat# ab120651, CAS No 93957-55-2), pravastatin (Sigma-Aldrich, Cat# 524403, CAS No 81131-70-6), pitavastatin (Sigma Aldrich, Cat# SML2473, CAS No 147526-32-7), benproperine (Hycultech, Cat# HY114657A, CAS No 19428-14-9), haloperidol (Sigma Aldrich, Cat# H1512, CAS No 52-86-8), CK666 (Sigma Aldrich, Cat# SML0006, CAS No 442633-00-3), CK869 (Sigma Aldrich, Cat# C9124, CAS No 388592-44-7). All compounds were dissolved in DMSO (AppliChem, Cat# A3672, CAS No 67-68-5). The final concentration of DMSO for all compounds and vehicle control was at maximum 0.1% and serially diluted down in lower concentrations, except for pravastatin, where it was 1%, indicated by “DMSO high” (included as additional experimental vehicle control).

### Assessment of *in vitro* viability

ASPC1 cells were seeded in triplicate per condition on 96-well plates (8 × 10^3^ to 20 × 10^3^ cells/well, depending on the time point). Compounds were added to ASCP1 cells at indicated concentrations. After 0, 24, 48 and 72 hours, cell viability was assessed using the PrestoBlue cell viability assay (Invitrogen, Cat #A13262) per manufacturer’s instructions (Supplementary Figure S1). Absorbance at 570 and 600 nm was measured with a Tecan Spark 10M plate reader (Tecan Trading AG, RRID:SCR_021897).

### Live-cell imaging

96-well plates were coated with Collagen I (Sigma Aldrich, Cat #C8919, CAS No 9007-34-5) for 1 hour (at 37 ℃ and 5% CO_2_), then washed once with PBS. Cells were seeded at 2,000 cells per well and incubated overnight. Cells were then treated with DMSO vehicle or respective compound for 48 hours prior to imaging. On each plate, four technical replicates per condition were imaged.

Time-lapse brightfield imaging was performed using a cell imaging multi-mode reader (Cytation 5, Agilent Technologies, RRID:SCR_019732). Each plate was imaged with the 20 × objective for 4 hours at 37 ℃ and 5% CO_2_ with a fixed number of kinetic reads/looping times as indicated. Individual images were aligned and stitched into .mp4 files using Gen5 software (RRID:SCR_017317). Videos of individual wells were excluded from further analysis if they were improperly aligned by Gen5 software or cell debris was misidentified as migrating cells.

### Computational segmentation and tracking

The tracking workflow was established as previously described (https://doi.org/-10.1038/s41467-023-43765-3). The generated movies, exported as .mp4 files, underwent automated segmentation using the cellular segmentation tool Cellpose (https://doi.org/10.1038/s41592-020-01018-x). To adapt to the domain-specific characteristics of the ASPC1 cell line, a pretrained Cellpose model was fine-tuned using representative samples.

Subsequent single-cell tracking was carried out using a classical tracking algorithm based on the Earth Mover’s Distance (https://doi.org/10.1007/978-3-319-10470-6_15). The tracking data contained unique tracking labels for each cell within the segmented images of the time-lapse sequence. Only cells in at least 10 consecutive frames were analyzed and reported as mean velocity pixel per frame. For each well, approximately 50–200 cells were measured in this way.

### Computational processing of cell migration data

Cell migration data were processed using the programming language R (R Core Team, 2020), resulting in a table with 30,037 rows (individual cells) and multiple columns representing cell-specific features, including:

- mean cell velocity *y*_*i*_ ∈ ℝ^+^
- well identifier, indexed as *w*[*i*] ∈ {1, 2, …, 346}
- plate identifier, indexed as *p*[*w*] ∈ {1, 2, …, 8}
- wells on plates *p* ∈ {1, 2, 3 } (generated as part of experiment A) and *p* ∈ { 4, 5, 6, 7, 8 } (generated as part of experiment B) were each treated with the same sets of 13 treatments; in total, there were 20 distinct treatments, indexed by *t* ∈ { 1, 2, …, 20 }, with only three treatments common to both plate groups
- each treatment represents a combination of a chemical compound and its administered concentration, applied across 4 wells per plate (referred to as technical replicates)

We observed between 10 and 275 cells per well (mean = 87 cells). Cell velocities ranged from 0.43 to 30.75 pixels/frame (mean velocity = 6.29 pixels/frame). Velocities on plates *p* ∈ { 1, 2, 3} were systematically lower compared to those on *p* ∈ {4, 5, 6, 7, 8 }, attributable to slight differences in imaging configurations used for the two plate groups. This batch effect is corrected by our model, as described in the following sections.

### Statistical modeling of cell migration velocity

We developed a Bayesian hierarchical regression model (*M*) to estimate treatment effects on cell migration velocity while accounting for experimental structure (technical/biological replicates, plate effects) and pooling information across groups.

This Bayesian approach is advantageous because it allows use of: (i) prior information (e.g. knowledge of effect magnitudes), (ii) explicit modeling of the experiment-specific structure (e.g. nested biological and technical replicates) and (iii) partial pooling of information across groups (Gelman et al., 2013).

#### Model structure

The observed migration velocity *y*_*i*_ for cell *i* was modeled using a gamma likelihood. The gamma distribution was chosen for the likelihood due to its flexibility in modeling positive, right-skewed data:

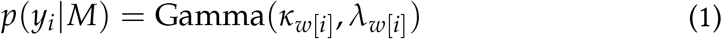

where *w*[*i*] is the index of well of cell *i*, and *κ*_*w*[*i*]_ ∈ ℝ^+^ and *λ*_*w*[*i*]_ ∈ ℝ^+^ are the shape and rate parameters of that cell. The mean (*µ*_*w*[*i*]_ ∈ ℝ^+^) of the gamma distribution is the ratio of *κ*_*w*[*i*]_ to *λ*_*w*[*i*]_, which allows reparameterization of the likelihood in terms of *κ*_*w*[*i*]_ and *µ*_*w*[*i*]_ as:

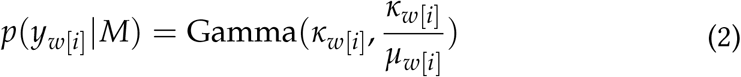

#### Accounting for experimental design

Each well *w* belongs to a plate *p* ∈ {1, 2, …, *P*} (biological replicate) and is treated by a treatment *t* ∈ {1, 2, …, *T*}. Multiple wells per plate may receive the same treatment (technical replicates, indexed as *pt*[*w*]). The log-transformed means of wells receiving the same treatment are modeled hierarchically, by treating the corresponding coefficients (log(*µ*_*pt*[*w*]_)) as random samples drawn from normal distributions:

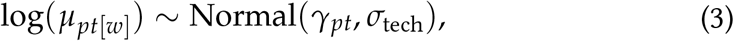

where *γ*_*pt*_ is mean effect of treatment *t* on plate *p*; and *σ*_tech_ captures the variability between technical replicates. However, we know that cell migration assays are affected by plate-specific batch effects that can systematically bias the migration velocity across all wells on the plates. To correct for plate-specific batch effects, we include an offset term *α*_*p*_, defined as the mean migration velocity (on log-scale) of a control treatment (e.g., DMSO) on plate *p*, effectively normalizing batch effects across plates:

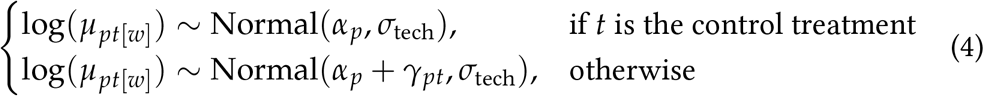

where the treatment effect *γ*_*pt*_ captures deviations from this baseline for treatment *t* on plate *p*; and *σ*_tech_ captures the variability between technical replicates.

Wells treated with the same treatment (*t*) will have similar cell migration velocities (similar *γ*_*pt*_ coefficients) after accounting for batch effects. Hence, the model treats *γ*_*pt*_ as random samples drawn from normal distributions:

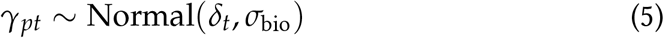

where δ_*t*_ is the overall (across plates) effects of treatment *t* on cell migration; and *σ*_bio_ is the standard deviation between biological replicates. The hierarchical structure enables sharing of information across plates while preserving treatment-specific estimates.

We observed log-linear relationship between the means and variances of the gamma distributed velocities across wells (Supplementary Figure S2A), which is expected from the linear relationship between the mean (*µ*_*w*_ = *κ*_*w*_/*λ*_*w*_) and variance 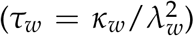 of the gamma distribution. The mean of the ratio, *µ*_*w*_/*τ*_*w*_, which is also the shape (*λ*_*w*_) of the gamma distribution, was close to 0.7 (Supplementary Figure S2B). Using the relationship between the shape (*λ*_*w*_) and rate (*κ*_*w*_) of the gamma distribution, we expect that *κ*_*w*_ has a similar magnitude as *µ*_*w*_:

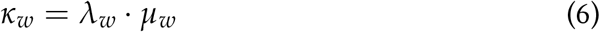

Hence, we assumed that values log(*κ*_*w*_) are also approximately normally distributed, and treated them as random samples drawn from a normal distribution with mean *µ*_*κ*_ and standard deviation *σ*_*κ*_:

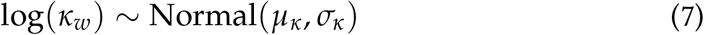

#### Quantifying treatment effects

For treatments *i* and *j*, the contrast *ρ*_*ij*_ = δ_*i*_ − δ_*j*_ represents the log-fold change in migration velocity. If *ρ <* 0 and the 95% Highest Density Intervals (HDIs) of *ρ* lie mostly or completely below 0 (0 = null effect), we have strong evidence of reduced cell velocity in *i* relative to *j*. On the other hand, if *ρ* and the 95% HDI of *ρ* lie mostly or completely above 0, we have strong evidence of increased cell velocity in *i* compared to *j*. Distributions with the 95% HDIs more or less centered around 0 indicate that there is no evidence for a clear change in the cell velocity. Note that unclear evidence is not equivalent to no change, because for a treatment group with *ρ*≈ 0 we may also have a wide 95% HDI, including possibilities for positive or negative change. cellmig outputs mean and 95% HDI of *ρ*, including the probability of differential effect on migration (*π*):

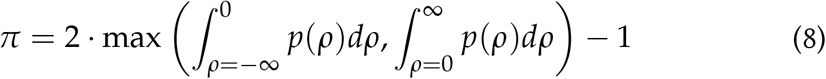

Here, *π* ≈ 1 indicates strong evidence for a difference in cell migration velocity between the treatments, while *π* ≈ 0 suggests negligible evidence.

For easier interpretation, cellmig exponentiates the log-fold-changes described by δ_*t*_, *γ*_*pt*_, and *ρ*_*ij*_, yielding fold-changes 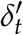, 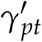, and 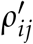, respectively.

#### Model priors

Our Bayesian approach requires prior distributions for the parameters *α*_*p*_, δ_*t*_, *σ*_tech_, *σ*_bio_, *µ*_*κ*_, and *σ*_*κ*_. We employed weakly informative priors that constrain parameters to plausible ranges while allowing the data to dominate posterior inferences. Using prior predictive checks, we ensured that the prior predictive distribution for the data has at least some density around extreme but plausible velocities (Supplementary Figure S3).

To assign a prior on *α*_*p*_, we evaluated the log-transformed cell velocities in our dataset, which were approximately normally distributed with mean *µ* ≈ 1.7 and standard deviation *σ* ≈ 0.5 (Supplementary Figure S2C). The well-specific mean velocities (on log-scale) were naturally lying within the distributions of the cell-specific velocities in both datasets. The parameter *α*_*p*_ represents the mean migration velocity of a control treatment (e.g., DMSO) on plate *p*, which we assume is also covered by the distribution of observed cell-specific velocities. To account for additional heterogeneity, we assign a prior on *α*_*p*_ which fully encloses the observed distribution of log-transformed cell velocities Supplementary Figure S2C):

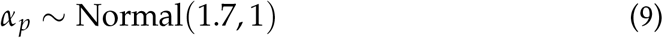

Based on our knowledge about the similar magnitudes of *µ*_*w*_ and *κ*_*w*_, we assumed that the overall mean of the population of *κ*_*w*_ (*µ*_*κ*_) should be similar to the overall mean of the population of *µ*_*w*_. Therefore, we assigned the same prior distribution to *µ*_*κ*_:

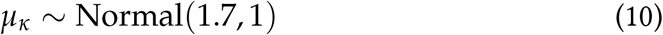

Meanwhile, δ_*t*_ represents the overall effect (generalized deflection from *α*_*p*_) on cell migration associated with treatment group *t*. If we assume δ_*t*_ = 1 and ignore the technical and biological variability in Equation 4 and 5, then from Equation 4 it follows that *µ*_*pt*[*w*]_ = exp(*α*_*p*_) · exp(1), i.e. the mean cell velocity increases by a factor of exp(1) = 2.71 over the baseline cell migration velocity of the plate. However, if δ_*t*_ = 2, then the mean cell velocity would increase by a staggering factor of exp(2) = 7.38 over the baseline migration. Using the control treatments in our experiments, we observed δ_*t*_=-0.52 (95% HDI=[-0.45, -0.59]) for the compound CK666 at 40 µM, and δ_*t*_=0.56 (95% HDI=[0.49, 0.64]) for fluvastatin at 10 µM, which are associated with severe reduction and increase in the cell migration velocity relative to DMSO (control treatment), respectively. Hence, for most treatments we expected |δ_*t*_| *<* 1, which implied that we need a prior distribution that assigns high density to values around 0, and low density to values larger than 1 and smaller than − 1. In fact, even if CK666 or fluvastatin were chosen as the the control treatment, |δ_*t*_| barely crosses the magnitude 1. Nevertheless, to account for additional heterogeneity (e.g. from newly synthesized chemical compounds with stronger effect on cell migration), we assigned a normal distribution as prior on δ_*t*_, with mean *µ* = 0 and standard deviation *σ* = 1, which assigns 95% of the density to values between −1.95 and 1.95:

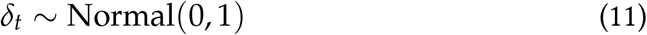

We used regularizing priors on the remaining scale parameters:

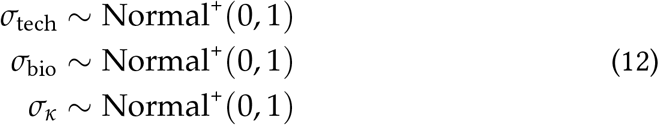

### Experiment planning by simulation

Our Bayesian model (*M*) is generative and can simulate cell velocities using the same error model (Equation 1). In this framework, the values of *κ*_*w*[*i*]_ and *λ*_*w*[*i*]_ are drawn from their priors, which are themselves informed by parameters in subsequent layers of the model. This simulation process is critical for optimizing experimental design, particularly in determining the optimal number of (i) cells, (ii) technical replicates, and (iii) biological replicates. These factors directly affect the uncertainty in estimating treatment effects (δ_*t*_) and their differences (*ρ*_*ij*_). The model can operate in two modes: *fully* generative, where all free parameters are sampled from their priors, or *partially* generative, where certain parameters are fixed by the user (e.g., to align with estimates from prior experiments).

## Results

### Experimental design and computational workflow

To evaluate the performance of cellmig in quantifying treatment effects on cell migration velocity, additionally taking into account multiple experimental variables such as technical/biological replicates, plate variability, we conducted two independent high-throughput experiments (A and B) using pancreatic cancer cells ASPC1 (Figure 1). This cell line was chosen as it exhibits robust migratory activity in live-cell assays (Dorsch et al., 2021). Experiments A and B comprised three and five biological replicates, respectively. Each well of the 96-well plate was treated with one of eight compounds at specified concentrations. Each condition was tested in four technical replicates per plate and in at least three independent experiments. While experiments A and B tested largely distinct compound sets, both included the vehicle-only control (DMSO) on all plates at the highest DMSO concentration of the compound-treated conditions. Additional controls included a migration enhancer (fluvastatin 10 µM) and reducer (ARP2/3 complex inhibitor CK666 40 µM). Both enhancer and reducer were included to implement upper and lower baselines to be used for cross-plate comparison.

**Figure 1:**
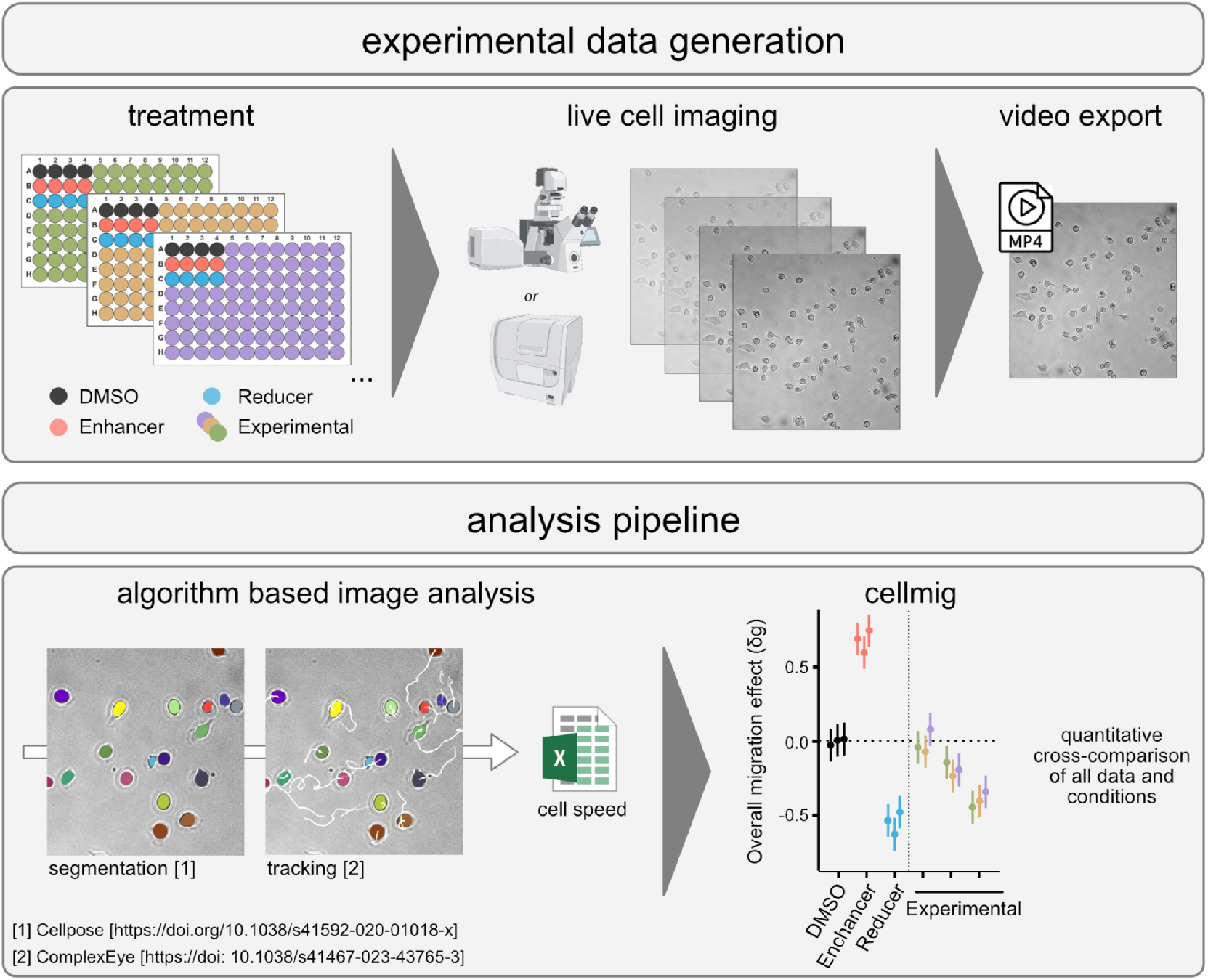
(Top) Live-cell imaging data is collected from multiple 96-well plates, each treated with matching controls (neutral, reducer, enhancer of migration) and unique experimental compounds in independent biological experiments. Still-frame images are captured at set intervals and stitched into a single video per well. (Bottom) To quantify single-cell migration velocity, videos undergo segmentation and tracking. The average velocity of each cell is calculated and exported as a spreadsheet for all experiments. cellmig enables quantification and cross-comparison of conditions using matching controls across plates.

In experiment A, experimental compounds included three statins, pitavastatin, fluvastatin, and pravastatin, each tested at 2.5, 5, and 10 µM, to allow for direct dose-response comparison. In this experiment, pitavastatin has a low solubility in DMSO. To therefore account for the higher concentrations of DMSO, “DMSO high” was included as an additional vehicle control setting. Experiment B included ARP2/3 inhibitors and antipsychotics, two classes of compounds reported to reduce cellular migration due to altered actin cytoskeleton dynamics (Chen et al., 2016). ARP2/3 inhibitors, CK869, CK666 (Ilatovskaya et al., 2013) and benoproperine (Yoon et al., 2019). These compounds were administered at 5 and 10 µM, each. For all tested compounds and concentrations, cell viability assays were performed to exclude potential bias on migration effects due to cell death (Supplementary Figure S1).

The subsequent computational analysis included movie generation, data quality control, cell segmentation, and cell tracking, as described in the Methods section. Together, experiments A and B generated a structured dataset representative of high-throughput cell migration assays. The dataset was exported as a spreadsheet, describing the observed velocity of *n* cells (rows) and their experimental features (chemical compound, dose, plate identifier, well identifier), which was used as main input by cellmig (version 0.99.14) to fit a hierarchical Bayesian model implemented in Stan (Carpenter et al., 2017). Using default parameters, cellmig performed model inference and validation (Supplementary Section S1). The posterior distributions of the model parameters were then used to quantify treatment effects, including their associated uncertainties, as detailed in the following sections.

### Overall treatment effects

Data was analyzed using cellmig, with DMSO as the reference for batch correction (Methods). The model reported overall treatment effects (δ_*t*_) as log-fold-changes relative to DMSO (Figure 2). Exponentiating δ_*t*_ yields interpretable fold-changes (vertical axis of Figure 2), and we refer to these estimates as 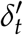, where 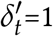 indicates no change in treatment *t* relative to DMSO.

**Figure 2:**
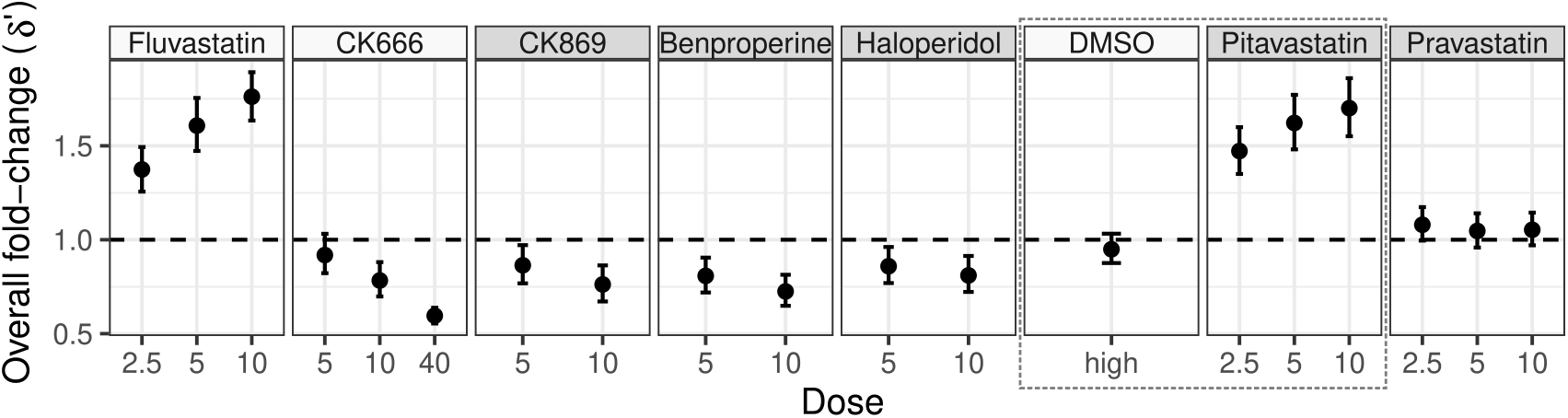
Overall treatment effects on cell migration velocity. Mean (dot) and 95% highest density interval (HDI; error bars) of the overall treatment effects (δ^*′*^), defined as combinations of compound and dose. Horizontal dashed line indicates the null effect (δ^*′*^=1). Control compounds (DMSO, CK666, and fluvastatin) have white panels. The dotted gray rectangle encloses results for pitavastatin (at each concentration) and the corresponding control (“DMSO high”).

As expected, high-dose fluvastatin (10 µM) and CK666 (40 µM) showed strong positive 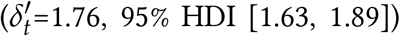 and negative 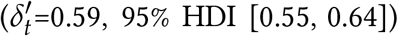 effects, respectively. Dose-response relationships were evident for control compounds: fluvastatin effects decreased to 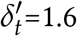, 95% HDI [1.47, 1.75] and 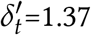, 95% HDI [1.25, 1.49] at 5 µM and 2.5 µM, respectively, while CK666 effects decreased to 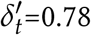, 95% HDI [0.7, 0.88] and 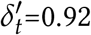, 95% HDI [0.82, 1.03] at 10 µM and 5 µM, respectively. High-dose DMSO caused a negligible velocity reduction 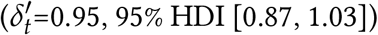 with 95% HDIs overlapping the null effect (δ^*′*^=1).

Among the compounds, pitavastatin (2.5-10 µM) increased migration 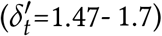, whereas CK869 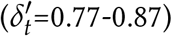, benproperine 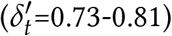, and haloperidol 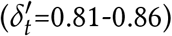 reduced migration at 5 and 10 µM. We observed dose-dependent responses in these treatment groups as well. In contrast, pravastatin showed negligible effects across all doses 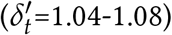 with 95% HDIs overlapping the null effect (δ^*′*^=1).

### Quantifying non-treatment contributions to variation

cellmig further quantified plate-specific treatment effects (*γ*_*pt*_ for treatment *t* on plate *p*; Figure 3B) relative to DMSO. As before, exponentiating *γ*_*pt*_ yielded interpretable fold-changes 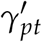 (vertical axis of Figure 3B). While 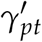 values generally mirrored the overarching 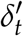 estimates, their variability across plates modeled by parameter *σ*_bio_ (Supplementary Figure S4B) accounted for biological heterogeneity in treatment responses. To assess batch correction efficacy, we compared 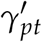 distributions for shared treatments (e.g., fluvastatin at 10 µM) against raw plate-specific mean velocities (Figure 3C). Despite systematic baseline differences between experiments – cells in experiment A exhibited higher raw velocities than those in experiment B (Figure 3D: blue vs. orange dots) – 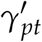 coefficients for shared treatments converged to similar values after correction (Figure 3B: blue vs. orange dots). In fact, the model parameters *α*_*p*_ accurately captured these inter-experiment biases (Supplementary Figure S4A): baseline velocities under DMSO were about 5-6 pixels/frame (experiment A) vs. 6-8 pixels/frame (experiment B).

**Figure 3:**
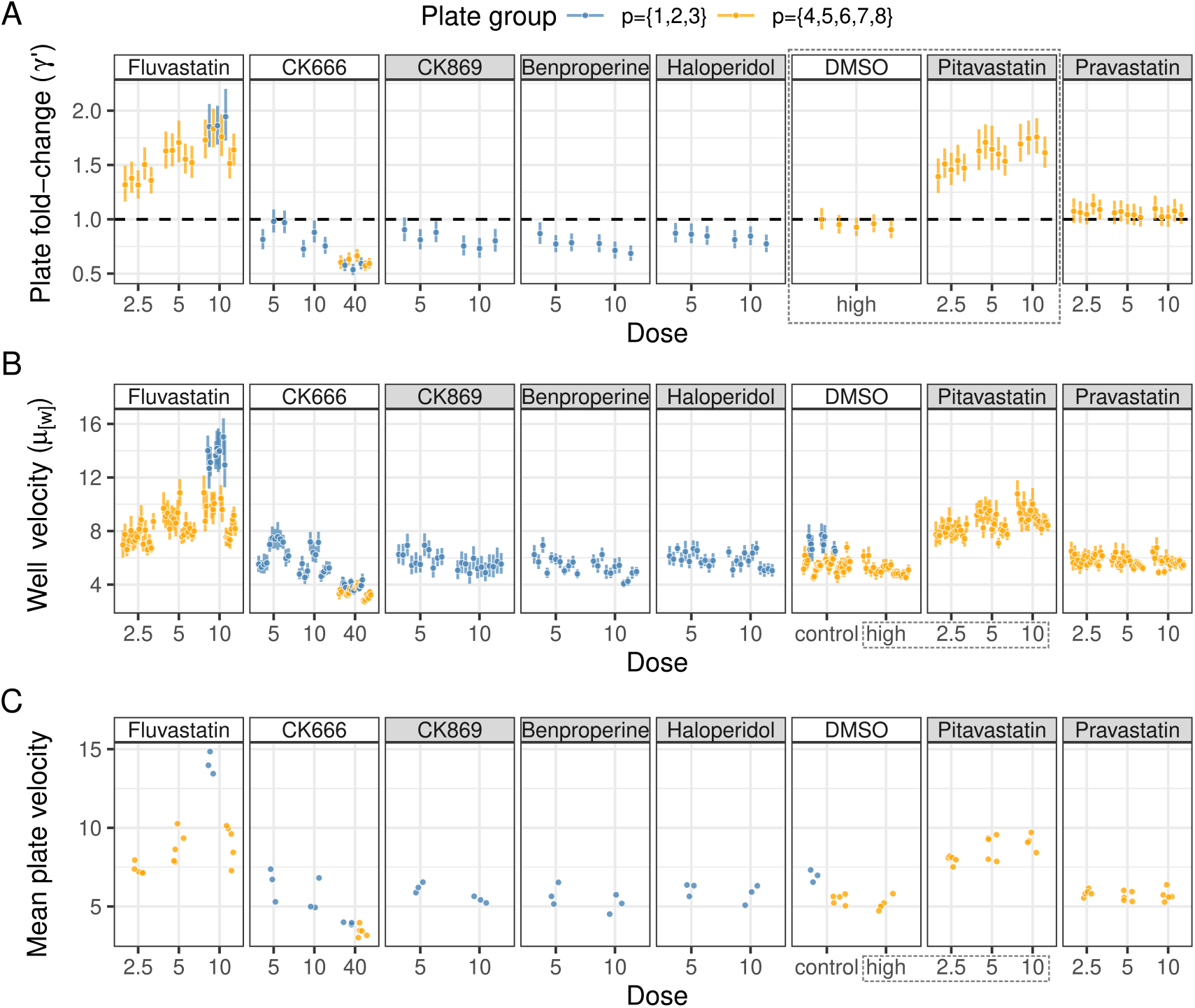
Observations and multilevel treatment effects on cell migration velocity. (A) Mean (dot) and 95% highest density interval (HDI; error bars) of the plate-specific treatment effects (*γ*^*′*^). (B) Inferred mean (dot) and 95% HDI (error bars) of the cell migration velocity in individual wells (*µ*_[*w*]_) before correction for plate-effects. (C) Observed mean (over all cells) treatment-specific velocities on each plate. Blue and orange dots and error bars in panels A-C correspond to plates from two different plate groups. Horizontal dashed line indicates the null effect (*γ*^*′*^=1). Control compounds (DMSO, CK666, and fluvastatin) have white panels. The dotted gray rectangles mark pitavastatin (at each concentration) and the corresponding control (“DMSO high”).

Using the lowest-level model parameters *µ*_*pt*[*w*]_, representing uncorrected well-specific mean velocities, we quantified technical variability across replicates. The distribution of *µ*_*pt*[*w*]_ estimates and their 95% HDIs revealed well-to-well noise, modeled by parameter *σ*_tech_ (Supplementary Figure S4B). This granular quantification enables not only accurate evaluation of treatment effects on migration velocities but it also facilitates the design of cell-migration experiments, as demonstrated below.

### Quantitative comparison of treatment effects

Our model quantifies treatment effects (δ_*t*_) relative to DMSO controls while enabling direct inter-treatment comparisons (e.g., between treatments *i* and *j*) through two key metrics: (1) fold change coefficients 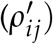; (2) probabilities (*π*_*ij*_) quantifying evidence for differential treatment effects.

Notably, pitavastatin (2.5-10 µM) increased migration by *ρ*^*′*^=1.7-2.3-fold compared to benproperine, CK869, and haloperidol (5-10 µM; red tiles, Figure 4A). These comparisons showed maximal probability (*π*=1; gray tiles, Figure 4B), indicating strong evidence for differential effects. In contrast, pravastatin (2.5-10 µM) exhibited weaker increase (*ρ*^*′*^=1.2-1.5-fold vs. suppressors; yellow tiles, Figure 4A), consistent with its negligible effect relative to DMSO. The velocity-suppressing compounds – benproperine, CK869, and haloperidol – showed comparable effects across doses (*ρ*^*′*^=0.83-1.06-fold) with limited evidence for differential activity (*π <* 1). As expected, comparisons of treatment groups with themselves yielded *ρ*^*′*^ = 1 and *π* = 0, i.e. no difference in treatment effects (diagonal entries, Figure 4A-B).

**Figure 4:**
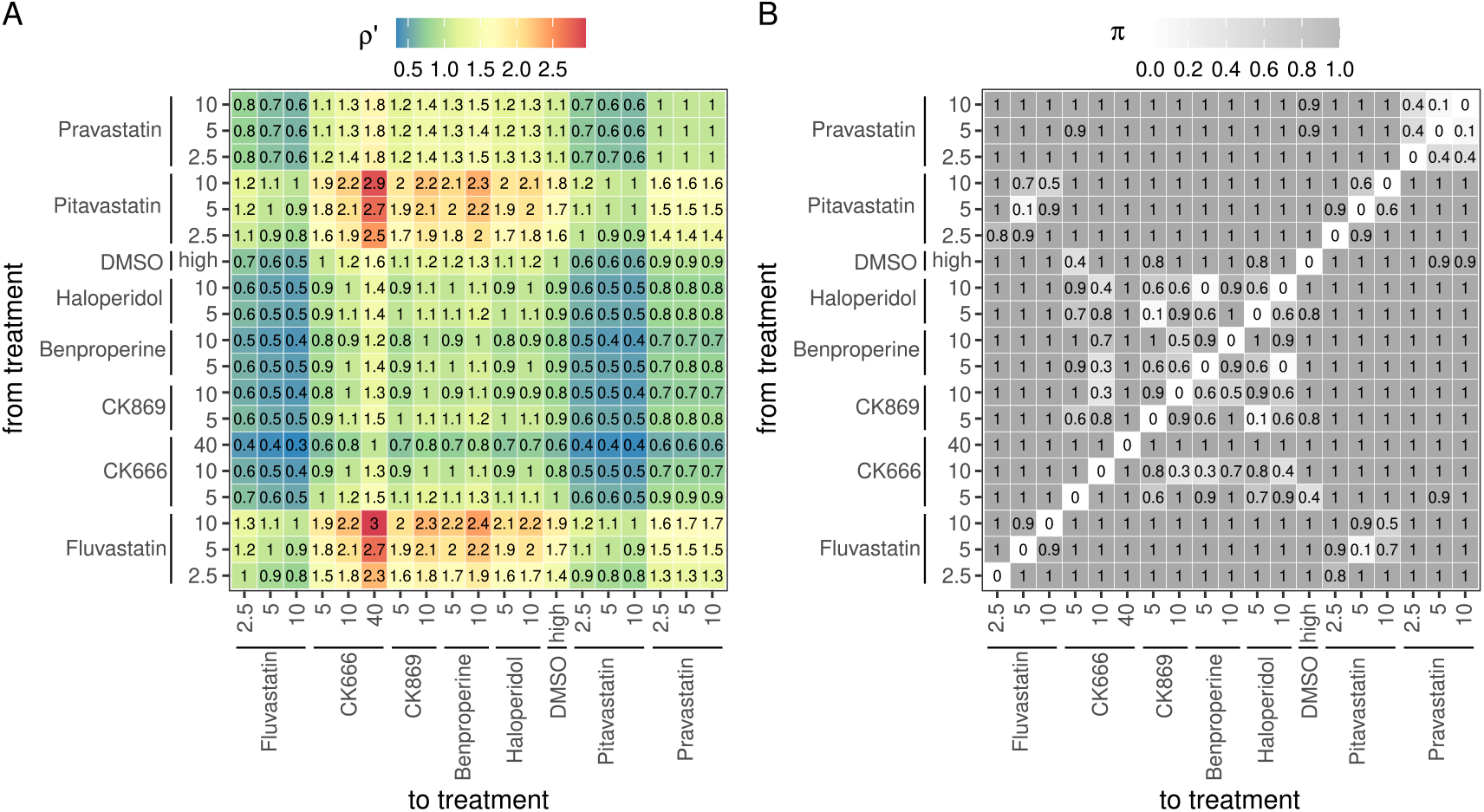
Differences in overall treatment effects. (A) Fold change 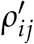 between overall treatments effects (δ_*t*_) of row (*i*) vs. column (*j*) treatment groups. Tile colors and labels represent 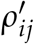 values (rounded to nearest tenth). (B) Probability of differential treatment effect described by parameter *π*_*ij*_. Tile colors and labels represent *π*_*ij*_ values (rounded to nearest tenth).

### Evaluating cellmig against state-of-the-art approaches

Cell migration data are most often analyzed with nonparametric tests (Cibir et al., 2023; Kleino et al., 2025; Masuzzo et al., 2013; Shannon et al., 2024; Wiggins et al., 2023; Wortel et al., 2021), such as the Wilcoxon rank-sum (Mann and Whitney, 1947) for pairwise, and the Kruskal-Wallis (Kruskal and Wallis, 1952) for multi-group comparisons. Given our 19 treatment groups, we benchmarked cellmig against the Kruskal-Wallis test followed by Dunn’s post hoc z-scores and *p*-values for 171 pairwise comparisons, with Benjamini-Hochberg FDR correction. We refer to this entire workflow as the H-test.

To mimic common workflows assuming minimal variability and negligible batch effects, we pooled cells by treatment, inflating sample sizes but ignoring experimental hierarchy–an issue avoided by cellmig. This revealed systematic discrepancies between H-test outputs (*p*-values, *z*-scores) and cellmig’s probabilistic metrics (*π, ρ*) (Figure 5).

**Figure 5:**
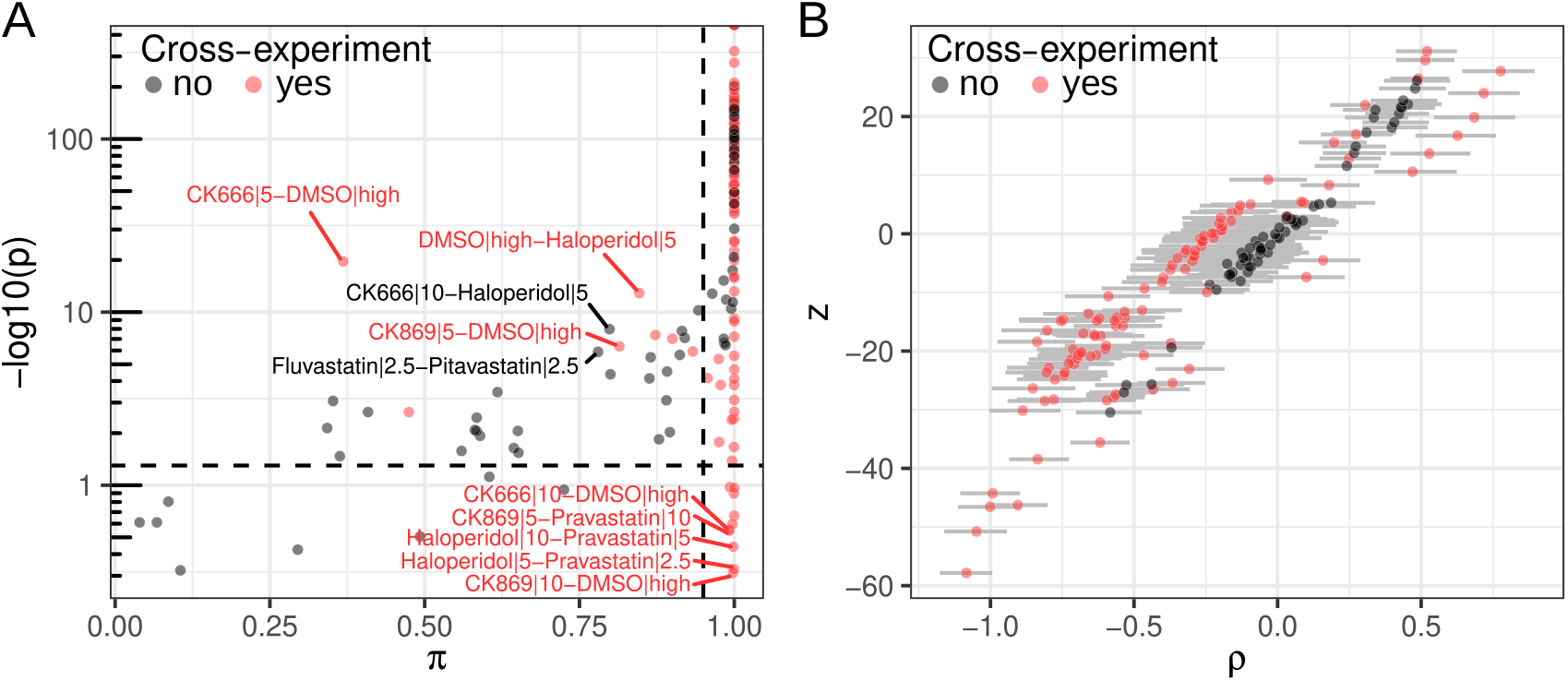
Benchmarking cellmig against the H-test. (A) Comparison of cellmig’s probabilities of differential treatment effects (*π*) and H-test’s (log_10_) adjusted *p*-values for 171 unique treatment pairs (dots). Labeled points highlight diverging results between cellmig and H-test. (B) Comparison of cellmig’s log-fold-change in treatment effect (*ρ*) and H-test’s *z*-scores for 171 unique treatment pairs (dots). Black/red dots denote intra-/inter-experiment pairs.

#### False negatives in H-test

Ten cross-experiment pairs (e.g., DMSO (high) vs. CK869 (10 µM) showed non-significant adjusted *p*-values (*p >* 0.05) in H-test despite strong evidence from cellmig (*π*≥ 0.95) (Figure 5A, bottom-right). All involved batch effects between Experiments A/B, where systematic velocity differences masked true treatment effects (Figure 3).

#### False Positives in H-Test

Fifteen pairs (e.g., CK666 (5 µM) vs. DMSO (high); CK666 (5 µM) vs. haloperidol (10 µM)) exhibited significant adjusted *p*-values (*p*≤ 0.05) but low *π* values (*π <* 0.95). While some false positives arose from unmodeled batch effects (e.g., cross-experiment pairs), others involved treatment groups from the same experiment. These discrepancies stemmed from two factors: (1) Pooling cells amplified apparent statistical power (green crosses vs. black dots in Supplementary Figure S5), rendering small effect sizes significant–a known challenge in large migration datasets (Shannon et al., 2024); (2) Applying the H-test to individual plates revealed fluctuating *p*-values (e.g., on log10-scale 0.86-3.53 for CK666 vs. haloperidol; 1-10.55 for fluvastatin vs. pitavastatin), reflecting non-negligible biological variability which is ignored by the H-test (Supplementary Figure S6). These inconsistent results, where some plates show significant *p*-values while others do not, highlight a key problem of the H-test, namely the interpretation of fluctuating *p*-values on individual plates.

In contrast, cellmig explicitly models biological variability, resulting in wider posterior distributions for overall treatment effects 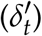. For example: CK666 (10 µM) vs. haloperidol (5 µM) had overlapping 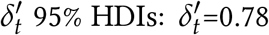, 95% HDI [0.82, 1.03] vs. 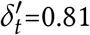, 95% HDI [0.72, 0.91]. Hence, the resulting *ρ*^*′*^=0.91, 95% HDI [0.78, 1.04] included 1 (the null effect) with *π*=0.8. We obtained similar results for fluvastatin (2.5 µM) vs. pitavastatin (2.5 µM).

#### Result robustness and interpretability

cellmig reports effect sizes 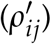 with uncertainties and evidence probabilities (*π*_*ij*_), while H-test relies on point estimates (*z*-scores, *p*-values). The *z*-score’s sensitivity to sample size amplifies trivial differences in large datasets (Figure 5B) and yields inconsistent interpretations across experiments with varying sample sizes. This fundamentally limits biological insight, as identical effect magnitudes may show divergent statistical significance. cellmig avoids this pitfall: its effect sizes remain stable across sample sizes, with the availability of more data improving precision rather than inflating significance.

### Experiment design by numerical simulation

Using similar parameter values similar to those inferred from our experimental data, including a synthetic vector of effect sizes 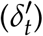 with values range from 0.5 to 2.0, we simulated cell velocity datasets while systematically varying three experimental dimensions: (1) *N*_cells_ = 25, 50, or 100 cells per well; (2) *N*_tech_ = 3, 6, 9, or 12 technical replicates; and (3) *N*_bio_ = 3, 6, 9, or 12 biological replicates per experiment. For each configuration (*N*_cells_, *N*_tech_, *N*_bio_), we generated 300 synthetic datasets and analyzed them using cellmig (Supplementary Section S2). We then evaluated the widths (*W*s) of the resulting 95% HDIs for 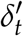 and evaluated their rate of overlap with synthetic effect sizes but not the null effect (true positive rate, TPR). Reliability of the *W* and TPR estimates was assessed by bootstrapping.

#### Biological replicates dominate uncertainty reduction

First we investigated the impact of experimental configuration (*N*_cells_, *N*_tech_, *N*_bio_) on *W* (Figure 6A). While increasing any of these three sample size dimension reduced *W*, the most substantial reduction occurred with increasing biological replicates (*N*_bio_).

**Figure 6:**
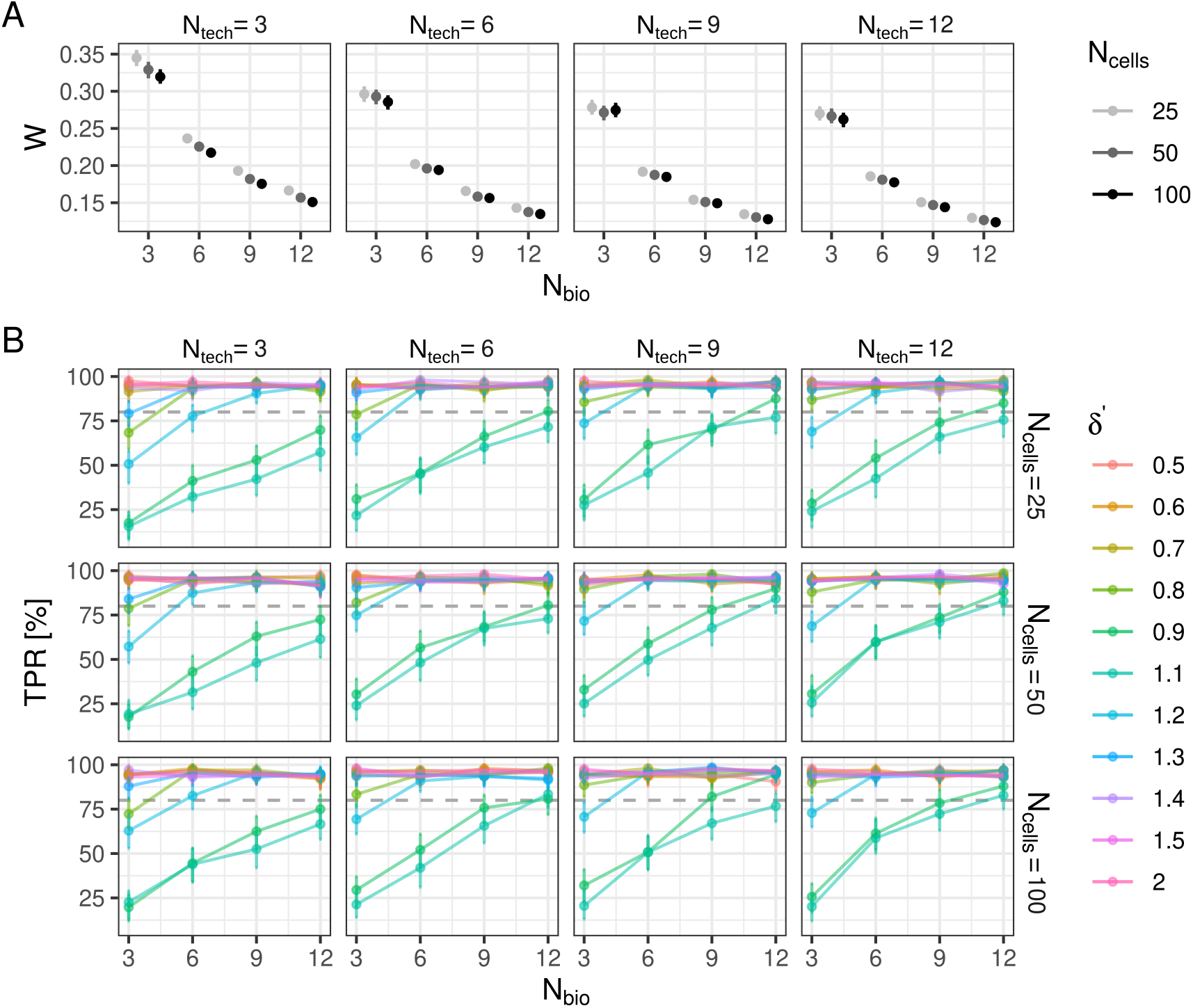
Power analysis. (A) Mean widths (*W*) of 95% Highest Density Intervals (HDIs) for effect sizes 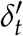 for different experimental configurations. Dots are mean *W* of 1,000 bootstraps. Error bars are 95% HDIs of bootstrapped*W*. Panels correspond to technical replicates (*N*_tech_). Colors represent cells per well (*N*_cells_), and x-axes show biological replicates (*N*_bio_). (B) True positive rates (TPR, %) for effect sizes 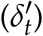. Lines connect TPR values (color-coded by 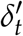) across *N*_bio_. Grid columns = *N*_tech_, rows = *N*_cells_. Dots are mean TPR values of 1,000 bootstraps, error bars are 95% HDIs of bootstrapped *TPR* values.

Specifically, setting *N*_bio_ to 3, 6, 9, and 12, was associated with consistently smaller *W* (Figure 6A) from *W*=0.35 to *W*=0.12, i.e., increasing *N*_bio_ to 6, 9, and 12 reduced *W* by factors of about 1.45, 1.8, and 2.1, respectively, relative to *N*_bio_ = 3 (Supplementary Figure S7A). Increasing *N*_tech_ shifted *W* to smaller values (panels in Figure 6A), however, we saw diminishing returns: *W* was reduced by a factor of 1.25 for *N*_tech_=12 vs. *N*_tech_=3 (Supplementary Figure S7B). Cell counts per well had negligible impact, with *N*_cells_=50 and 100 reducing *W* by less than 1.1-fold versus *N*_cells_=25 (Supplementary Figure S7C).

In summary, experimental designs should therefore prioritize biological replicates for minimizing uncertainty in 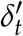. Technical replicates showed limited impact in these simulations, consistent with our assumption of low technical variability based on empirical data. We note that cellmig enables power analysis under diverse experimental conditions (e.g., high variability regimes), where technical replicates may have greater influence on *W*.

#### Controlling the number of true positives

A narrow uncertainty interval is only valuable if it actually contains the true effect size. We quantified whether an uncertainty interval contains the true effect size (here synthetic fold-changes 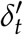), but not the null effect 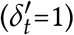. If it does, we call this a true positive. Otherwise, if the uncertainty interval contains the null effect, we call this a false negative. From 300 simulations performed with the same experimental configuration we computed the TPR (Figure 6B).

Large effects (e.g., migration enhancing 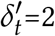, 1.5 and 1.4; or migration reducing 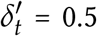, 0.6 and 0.7) achieved high TPR≥ 90% across all designs. Smaller effects (e.g., 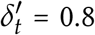 or 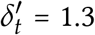 and 1.2) required at least 6 biological replicates for TPR ≥ 80%. For the smallest effects in our simulation 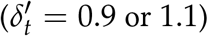, maximum number of biological replication (*N*_bio_ = 12) combined with at least 6 technical replicates yielded TPR close to 80%. This demonstrates that detecting subtle treatment effects (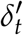 close to 1) requires substantial biological replication.

## Discussion

We have developed and validated a statistical model to rigorously quantify treatment-specific effects on cell migration velocity in high-throughput imaging assays. Our approach addresses key limitations of current methods by modeling hierarchical experimental designs and integrating sources of technical and biological variation into a unified Bayesian framework, implemented in the free, open source R package cellmig, with extensive vignettes for interdisciplinary researchers.

Existing analytical pipelines rely on non-parametric tests or descriptive summaries that fail to account for replicate structure, batch effects, or uncertainty propagation. Such methods risk overinterpreting single experiments, conflating technical noise with biological signals, and producing irreproducible results. In contrast, cellmig employs a Bayesian hierarchical model to deliver probabilistic estimates of condition-specific effects while explicitly accommodating nested data structures (e.g., plates, wells, cells). This enables statistically robust comparisons across conditions with quantified uncertainty, reducing susceptibility to batch artifacts and improving reproducibility.

The reproducibility of our findings underscores the robustness of the method. Compounds targeting ARP2/3 complex-mediated actin nucleation – CK666 and CK869 (Cao et al., 2024) – consistently reduced migratory capacity in a dose-dependent manner. Similarly, benproperine, an inhibitor of the ARPC2 subunit of the ARP2/3 complex, demonstrated a comparable dose-dependent reduction in migration, in line with prior observations (Yoon et al., 2019). Haloperidol, an antipsychotic drug previously shown to inhibit glioblastoma cell migration (Papadopoulos et al., 2020), also reduced migration in our screen, though to a lesser extent than the other compounds.

We previously reported that fluvastatin enhances pancreatic cancer cell migration by inducing a mesenchymal phenotype (Dorsch et al., 2021). Using cellmig, we not only confirmed its potent migration-inducing effect but also demonstrated a clear dose-dependent response mirrored by its analog, pitavastatin – both belonging to the same class of synthetic statins. In contrast, pravastatin, from the class of natural or fungal-derived derived statins, showed no effect on migration, suggesting that migration modulation may involve additional factors beyond HMG-CoA reductase inhibition.

Most importantly, cellmig enabled for the first time the direct and quantitative comparison of two completely independent experimental datasets, allowing for a rigorous assessment of compound-specific effects on migration and meaningful biological conclusions.

Another unique strength of cellmig is its generative capacity: the model can simulate synthetic datasets under user-defined assumptions, supporting power analysis and experimental design optimization. This feature is particularly valuable for resource-limited studies or large-scale screens, where balancing replicate numbers, batch configurations, and inferential precision is critical.

However, our approach has limitations. First, cellmig assumes data adhere to a predefined hierarchical structure (Supplementary Figure S8), which, while common in high-throughput imaging, may not fit alternative designs (e.g., simpler studies without technical replicates; or more complex studies including unmodeled confounders). Second, the current implementation focuses on mean migration velocity; while the framework is extensible to other phenotypes (e.g., persistence, directionality), such adaptations require additional methodological development. Third, the model priors are optimized for cell velocity expressed in pixels per frame. If velocity is given in other units, such as micrometers per second, the priors may need adjustment, which is supported in cellmig.

In conclusion, cellmig addresses a critical gap in the analysis of live-cell migration assays. By combining hierarchical Bayesian modeling with tools for simulation and uncertainty quantification, it enhances the rigor, interpretability, and cross-study comparability of high-throughput migration data – advancements essential for both mechanistic research and translational applications such as drug discovery.

## Acknowledgements

We would like to thank Eva Hahn and Nancy Meyer for technical assistance and members of the Grüner and Hoffmann groups for insightful discussions.

## References

Andy Aman and Tatjana Piotrowski. Cell migration during morphogenesis. Developmental biology, 341(1): 20–33, 2010.

LuYan Cao, Shaina Huang, Angika Basant, Miroslav Mladenov, and Michael Way. Ck-666 and ck-869 differentially inhibit arp2/3 iso-complexes. EMBO reports, 25(8): 3221–3239, 2024.

Bob Carpenter, Andrew Gelman, Matthew D Hoffman, Daniel Lee, Ben Goodrich, Michael Betancourt, Marcus Brubaker, Jiqiang Guo, Peter Li, and Allen Riddell. Stan: A probabilistic programming language. Journal of statistical software, 76 (1), 2017.

Mao-Liang Chen, Fu-Ming Tsai, Ming-Cheng Lee, and Yi-Yin Lin. Antipsychotic drugs induce cell cytoskeleton reorganization in glial and neuronal cells via rho/cdc42 signal pathway. Progress in Neuro-Psychopharmacology and Biological Psychiatry, 71: 14–26, 2016.

Zülal Cibir, Jacqueline Hassel, Justin Sonneck, Lennart Kowitz, Alexander Beer, Andreas Kraus, Gabriel Hallekamp, Martin Rosenkranz, Pascal Raffelberg, Sven Olfen, et al. Complexeye: a multi-lens array microscope for highthroughput embedded immune cell migration analysis. Nature Communications, 14(1): 8103, 2023.

Madeleine Dorsch, Manuela Kowalczyk, Mélanie Planque, Geronimo Heilmann, Sebastian Urban, Philip Dujardin, Jan Forster, Kristina Ueffing, Silke Nothdurft, Sebastian Oeck, et al. Statins affect cancer cell plasticity with distinct consequences for tumor progression and metastasis. Cell Reports, 37(8), 2021.

Peter Friedl and Darren Gilmour. Collective cell migration in morphogenesis, regeneration and cancer. Nature reviews Molecular cell biology, 10(7): 445–457, 2009.

Andrew Gelman, John B Carlin, Hal S Stern, David B Dunson, Aki Vehtari, and Donald B Rubin. Bayesian data analysis. Chapman and Hall/CRC, 2013.

Keren I Hulkower and Renee L Herber. Cell migration and invasion assays as tools for drug discovery. Pharmaceutics, 3(1): 107–124, 2011.

Daria V Ilatovskaya, Vladislav Chubinskiy-Nadezhdin, Tengis S Pavlov, Leonid S Shuyskiy, Viktor Tomilin, Oleg Palygin, Alexander Staruschenko, and Yuri A Negulyaev. Arp2/3 complex inhibitors adversely affect actin cytoskeleton remodeling in the cultured murine kidney collecting duct m-1 cells. Cell and tissue research, 354: 783–792, 2013.

Iivari Kleino, Mats Perk, António GG Sousa, Markus Linden, Julia Mathlin, Daniel Giesel, Paulina Frolovaite, Sami Pietilä, Sini Junttila, Tomi Suomi, et al. Cellromer: an r package for clustering cell migration phenotypes from microscopy data. Bioinformatics Advances, page vbaf069, 2025.

Nina Kramer, Angelika Walzl, Christine Unger, Margit Rosner, Georg Krupitza, Markus Hengstschläger, and Helmut Dolznig. In vitro cell migration and invasion assays. Mutation Research/Reviews in Mutation Research, 752(1): 10–24, 2013.

William H Kruskal and W Allen Wallis. Use of ranks in one-criterion variance analysis. Journal of the American statistical Association, 47(260): 583–621, 1952.

Rachel M Lee, Michele I Vitolo, Wolfgang Losert, and Stuart S Martin. Distinct roles of tumor associated mutations in collective cell migration. Scientific reports, 11(1): 10291, 2021.

Andrew D Luster, Ronen Alon, and Ulrich H Von Andrian. Immune cell migration in inflammation: present and future therapeutic targets. Nature immunology, 6(12): 1182–1190, 2005.

Henry B Mann and Donald R Whitney. On a test of whether one of two random variables is stochastically larger than the other. The annals of mathematical statistics, pages 50–60, 1947.

Paola Masuzzo, Niels Hulstaert, Lynn Huyck, Christophe Ampe, Marleen Van Troys, and Lennart Martens. Cellmissy: a tool for management, storage and analysis of cell migration data produced in wound healing-like assays. Bioinformatics, 29(20): 2661–2663, 2013.

Paola Masuzzo, Marleen Van Troys, Christophe Ampe, and Lennart Martens. Taking aim at moving targets in computational cell migration. Trends in cell biology, 26(2): 88–110, 2016.

Fotios Papadopoulos, Rafaela Isihou, George A Alexiou, Thomas Tsalios, Evrysthenis Vartholomatos, Georgios S Markopoulos, Chrissa Sioka, Pericles Tsekeris, Athanasios P Kyritsis, and Vasiliki Galani. Haloperidol induced cell cycle arrest and apoptosis in glioblastoma cells. Biomedicines, 8(12): 595, 2020.

R Core Team. R: A Language and Environment for Statistical Computing. R Foundation for Statistical Computing, Vienna, Austria, 2020. URL https://www.R-project.org/.

Germán Reig, Eduardo Pulgar, and Miguel L Concha. Cell migration: from tissue culture to embryos. Development, 141(10): 1999–2013, 2014.

Michael J Shannon, Shira E Eisman, Alan R Lowe, Tyler FW Sloan, and Emily M Mace. cellplato–an unsupervised method for identifying cell behaviour in heterogeneous cell trajectory data. Journal of Cell Science, 137(20), 2024.

Elvira Toscano, Elena Cimmino, Fabrizio A Pennacchio, Patrizia Riccio, Alessandro Poli, Yan-Jun Liu, Paolo Maiuri, Leandra Sepe, and Giovanni Paolella. Methods and computational tools to study eukaryotic cell migration in vitro. Frontiers in Cell and Developmental Biology, 12:1385991, 2024.

Laura Wiggins, Alice Lord, Killian L Murphy, Stuart E Lacy, Peter J O’Toole, William J Brackenbury, and Julie Wilson. The cellphe toolkit for cell phenotyping using time-lapse imaging and pattern recognition. Nature Communications, 14(1): 1854, 2023.

Inge MN Wortel, Annie Y Liu, Katharina Dannenberg, Jeffrey C Berry, Mark J Miller, and Johannes Textor. Celltrackr: an r package for fast and flexible analysis of immune cell migration data. ImmunoInformatics, 1:100003, 2021.

Yae Jin Yoon, Young-Min Han, Jiyeon Choi, Yu-Jin Lee, Jieun Yun, Su-Kyung Lee, Chang Woo Lee, Jong Soon Kang, Seung-Wook Chi, Jeong Hee Moon, et al. Benproperine, an arpc2 inhibitor, suppresses cancer cell migration and tumor metastasis. Biochemical pharmacology, 163: 46–59, 2019.

