## Supplementary for "Uncertainty-aware quantitative analysis of high-throughput live cell migration data"

This document includes:
- Supplementary Sections S1-S2
- Supplementary Figures S1-S10
- Supplementary References

### Supplementary Sections

#### Supplementary Section S1: Model inference and checks

Model inference in cellmig was performed using the No-U-Turn sampler in the rstan R package (version 2.32.7), with four Markov chains run for 2,000 iterations each, including 1,000 warm-up iterations. To assess model validity, we performed posterior predictive checks, which showed that the simulated data were consistent with the observed data (Supplementary Figures S9 and S10). We evaluated convergence using the potential scale reduction factor (PSRF), the effective sample size ($N_{\text{eff}}$), and diagnostic information provided by rstan (e.g., divergence warnings during MCMC sampling). After having passed these tests, the posterior distributions of the model parameters were used to quantify differences in treatment effects including their uncertainties.

#### Supplementary Section S2: Experiment design by numerical simulation

Based on our experimental data, we found that $\delta'_t$ values range from 0.5 (indicating a two-fold reduction in cell velocity for treatment group $t$ relative to control) to 2 (indicating a two-fold increase). Using these realistic parameter values, we generated a synthetic vector of treatment effects spanning small to large magnitudes: $\delta'_t$ = 0.5, 0.6, 0.7, 0.8, 0.9, 1.0, 1.1, 1.2, 1.3, 1.4, 1.5, 2.0, where the treatment group with index $t = 6$ ($\delta'_t = 1.0$) served as the control (offset). Using $\delta'_t$ and additional realistic model parameters from our experiments (e.g., biological replicate variability: $\sigma_{\text{bio}}$=0.1; technical replicate variability: $\sigma_{\text{tech}}$=0.1; Supplementary Figure S4), we simulated cell velocity datasets while systematically varying three experimental dimensions: (1) $N_{\text{cells}}$ = 25, 50, or 100 cells per well; (2) $N_{\text{tech}}$ = 3, 6, 9, or 12 wells (technical replicates) per plate; and (3) $N_{\text{bio}}$ = 3, 6, 9, or 12 plates (biological replicates) per experiment. For each configuration defined by the tuple ($N_{\text{cells}}$, $N_{\text{tech}}$, $N_{\text{bio}}$), we generated 300 synthetic datasets and analyzed them using cellmig.

We then evaluated the widths ($W$s) of the resulting 95% HDIs all $\delta'_t$ at a given experimental configuration, and evaluated their overlap status ($O$) with synthetic effect sizes. If the 95% HDI interval contains the true effect size (here synthetic fold-changes $\delta'_t$), but not the null effect ($\delta'_t$=1), we call this a true positive ($O$=1). Otherwise, we call this a false negative ($O$=0). From 300 simulations performed with the same experimental configuration we computed the true positive rate (percentage) as the mean of $O$: TPR = $\sum_{i=1}^{300}(O_i)/300 * 100$. Reliability of the $W$ and TPR estimates was assessed by the following bootstrapping procedure:

1. For each experimental configuration we performed 1,000 bootstrap iterations
2. In each iteration, we sample 100 $W$ and $O$ values with replacement for a specific experimental configuration and computed their means
3. This gives us for each experimental configuration a distribution of 1,000 mean $W$ and TPR values, that we characterize with its mean and 95% HDI.

### Supplementary Figures

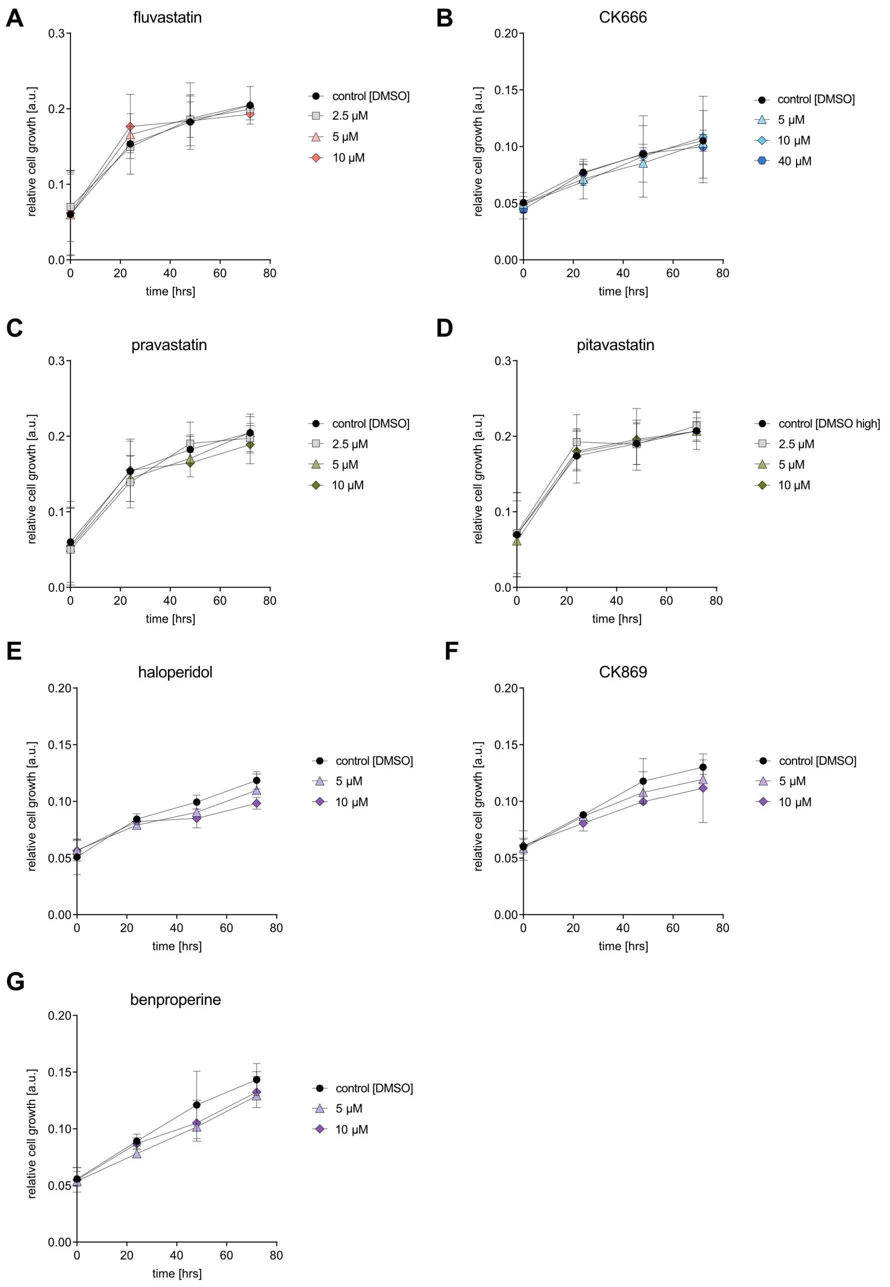


Supplementary Figure S1. Cell viability assay for (A) fluvastatin (B) CK666 (C) pravastatin (D) pitavastatin (E) haloperidol (F) CK869, and (G) benproperine in ASPC1 cells, tested at indicated concentrations and timepoints. DMSO at the concentration present in the highest compound concentration was used as vehicle only control. The error bars indicate +/- SEM.

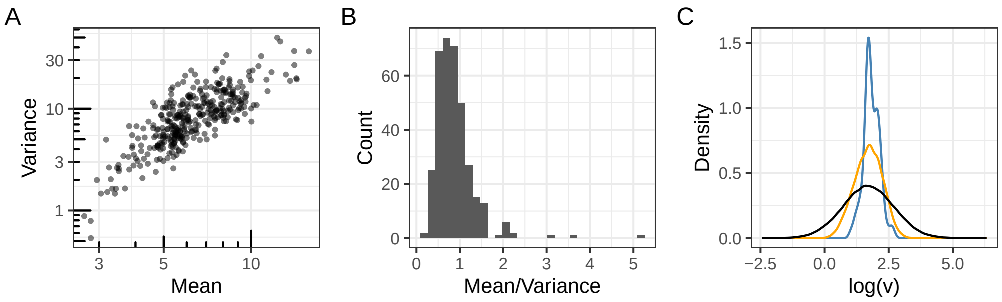


Supplementary Figure S2. Relationship between well-specific cell velocity means and variances. (A) Log-linear relationship between the variance (y-axis) and mean (x-axis) of well-specific cell velocities. (B) Ratio between well-specific cell velocity means and variances. (C) Probability distributions of individual cells (orange), well-specific means (blue), and the normal prior distribution (black) for $\alpha_p$ and $\mu_\kappa$, with mean $\mu = 1.7$ and standard deviation $\sigma = 1$. Panels were generated using the R package ggplot2 (v3.5.2) [Wickham, 2016] and combined with patchwork (v1.3.0) [Pedersen, 2025].

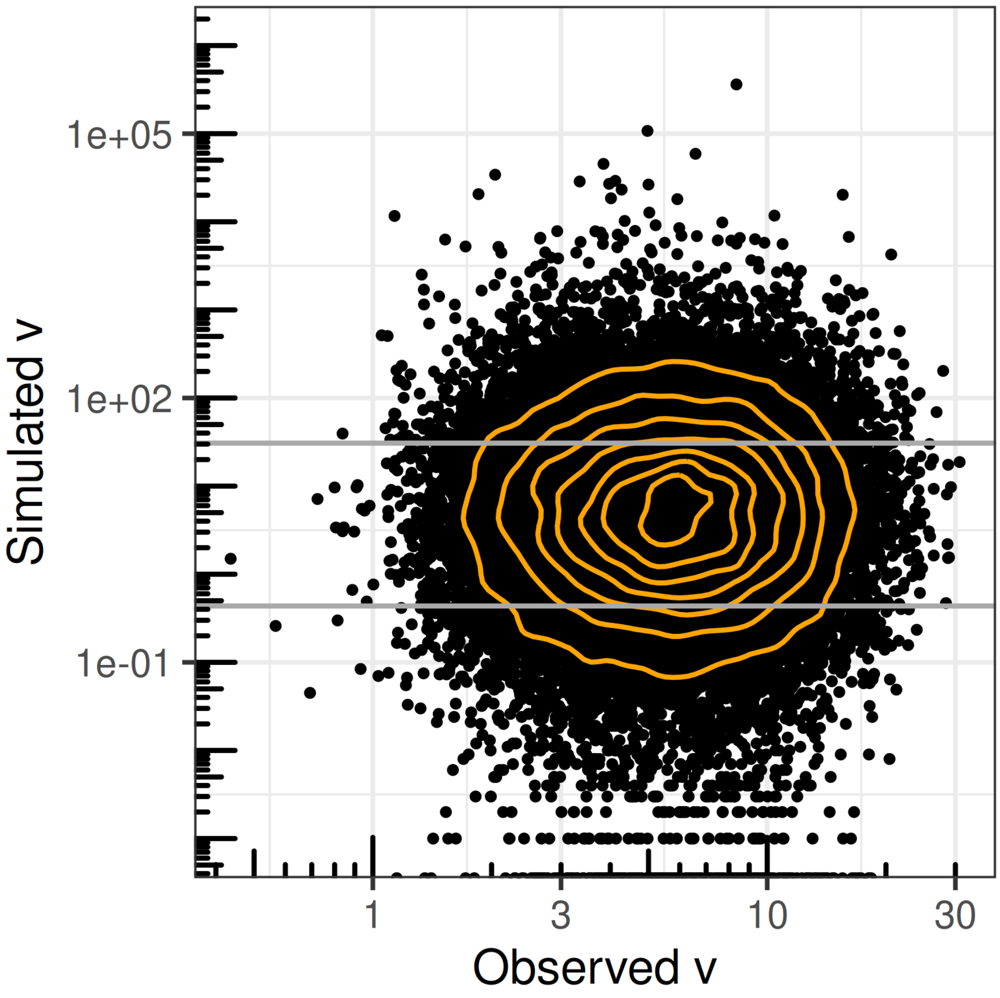


Supplementary Figure S3. Prior predictive distribution of cell velocities. Simulated and observed cell velocities ($v$) for 31,471 cells from dataset $D$ are plotted on the y-axis and x-axis, respectively, with both axes on a log10 scale. Simulated velocities of zero, are displayed along the bottom edges of the plots. Orange contours represent the two-dimensional density of the data. Gray horizontal lines indicate the minimum and maximum of the observed velocity distribution. The figure was generated using ggplot2 (v3.5.2) [Wickham, 2016].

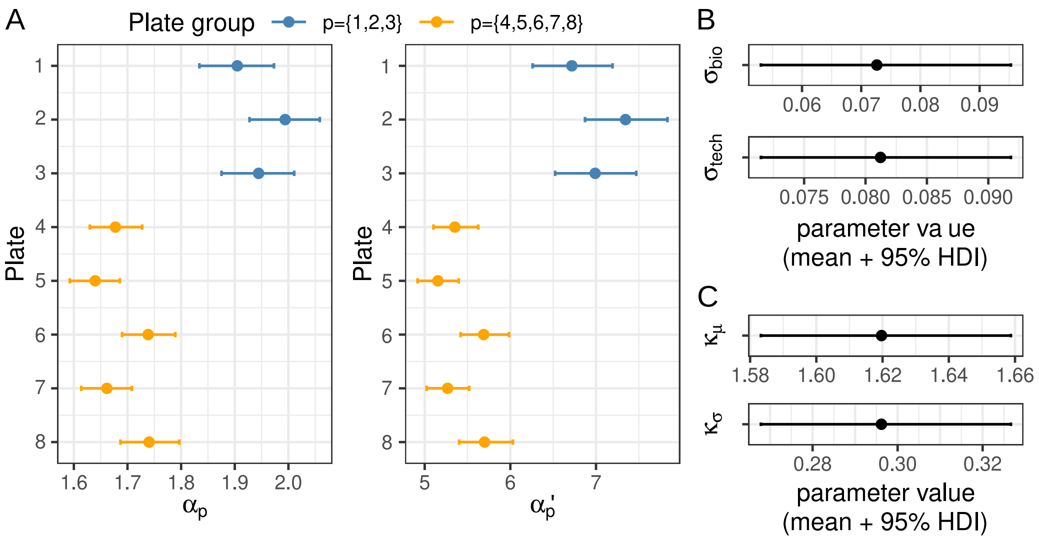


Supplementary Figure S4. Means and 95% HDIs of several model $M$ parameters. (A) $\alpha_p$ defined as the mean migration velocity (on log-scale) of the control treatment DMSO on each plate, and the exponentiated version, $\alpha'_p$, defined as the mean migration velocity (pixels/frame). Blue and orange dots and error bars correspond to plates from two different plate groups; (B) $\sigma_{bio}$ defined as the standard deviation of treatment effects between biological replicates (plates) and $\sigma_{tech}$ defined as the standard deviation of treatment effects between technical replicates (wells on plate). (C) mean ($\mu_\kappa$) and standard deviation ($\sigma_\kappa$) of the normal distribution of well-specific $\log(\kappa_w)$ parameters.

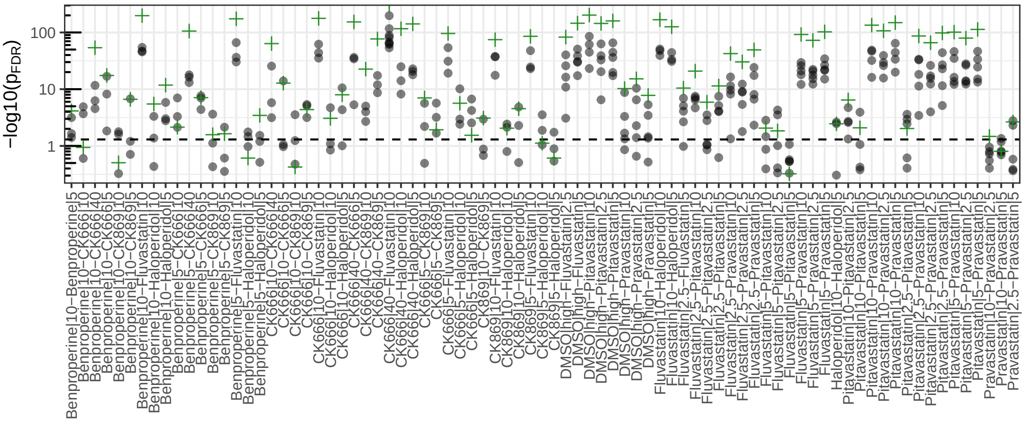


Supplementary Figure S5. Statistical comparison of cell velocity between pairs of treatment groups analyzed on the same plate. Black dots represent comparisons based on cell velocity data from individual plates: y-axis shows the $-\log_{10}$ $p$-values obtained from a post hoc analysis using the Dunn's test Benjamini-Hochberg FDR correction; x-axis shows the pairs of compared treatment groups. Green crosses represent the same statistical comparisons based on pooled cell velocities across plates. The horizontal dashed line represents $p$=0.05.

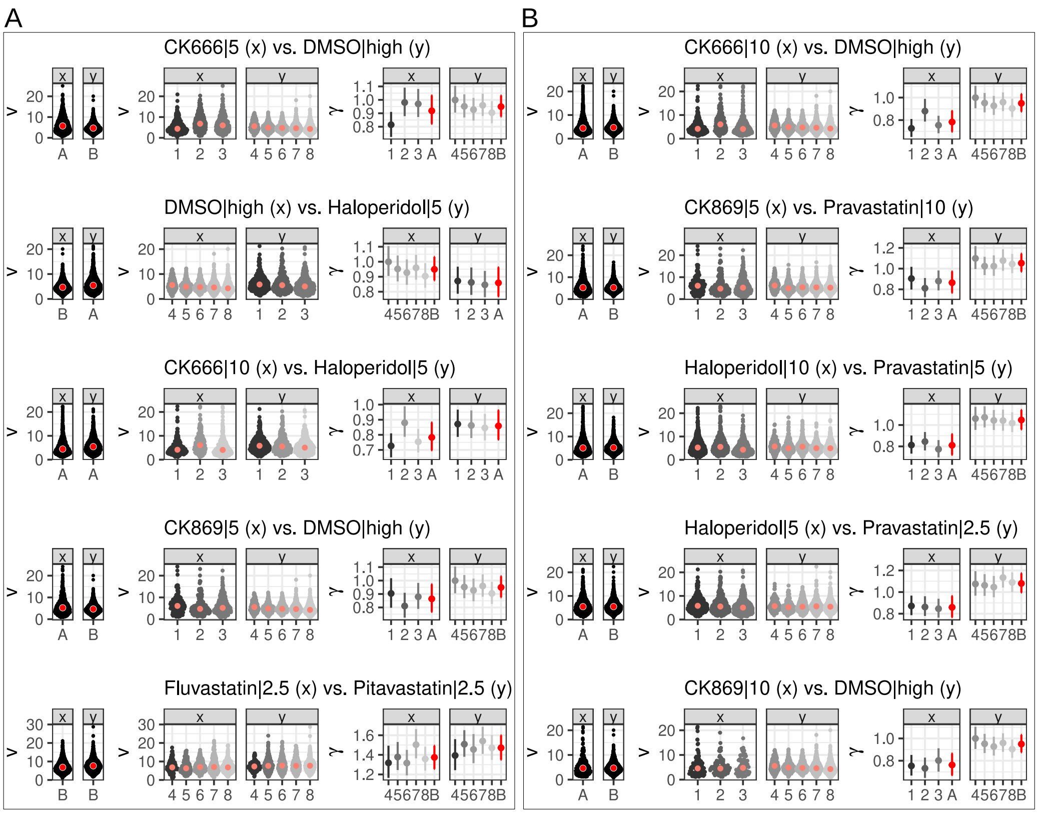


Supplementary Figure S6. (A) Five treatment pairs where the H-test yields false negatives. (B) Five pairs where the H-test yields false positives. Dots are cell velocities in each treatment group (x and y). The cells are shown as pooled across plates from experiment A and B (left panels), or within individual plates (middle panels). Mean plate-specific treatment effects and overall treatment effects and their 95% HDIs are shown as gray/red dots and error bars (right panels). Larger dots are group medians.

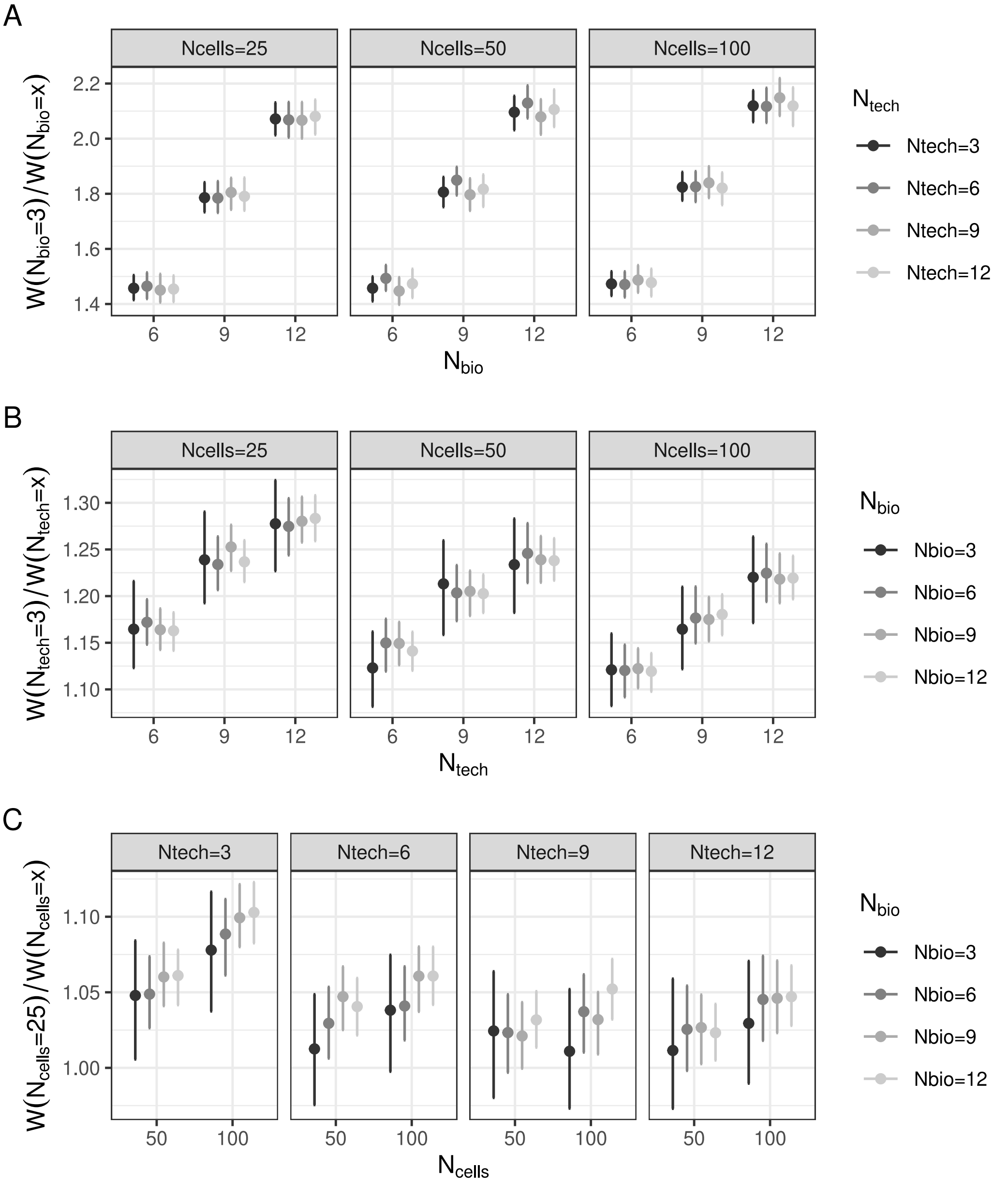


Supplementary Figure S7. Fold change in 95% HDI length ($W$) of $\delta_t'$ between pairs of experimental configurations. (A) Fold change in $W$ between $N_{\text{bio}}$=3 vs. $N_{\text{bio}}$=6, 9, and 12 for specific combinations of $N_{\text{tech}}$ and $N_{\text{cells}}$. (B) Fold change in $W$ between $N_{\text{tech}}$=3 vs. $N_{\text{tech}}$=6, 9, and 12 for specific combinations of $N_{\text{bio}}$ and $N_{\text{cells}}$. (C) Fold change in $W$ between $N_{\text{cells}}$=25 vs. $N_{\text{cells}}$=50 and 100 for specific combinations of $N_{\text{bio}}$ and $N_{\text{tech}}$. Dots are mean fold changes of 1,000 bootstraps, error bars are 95% HDIs of bootstrapped fold changes.

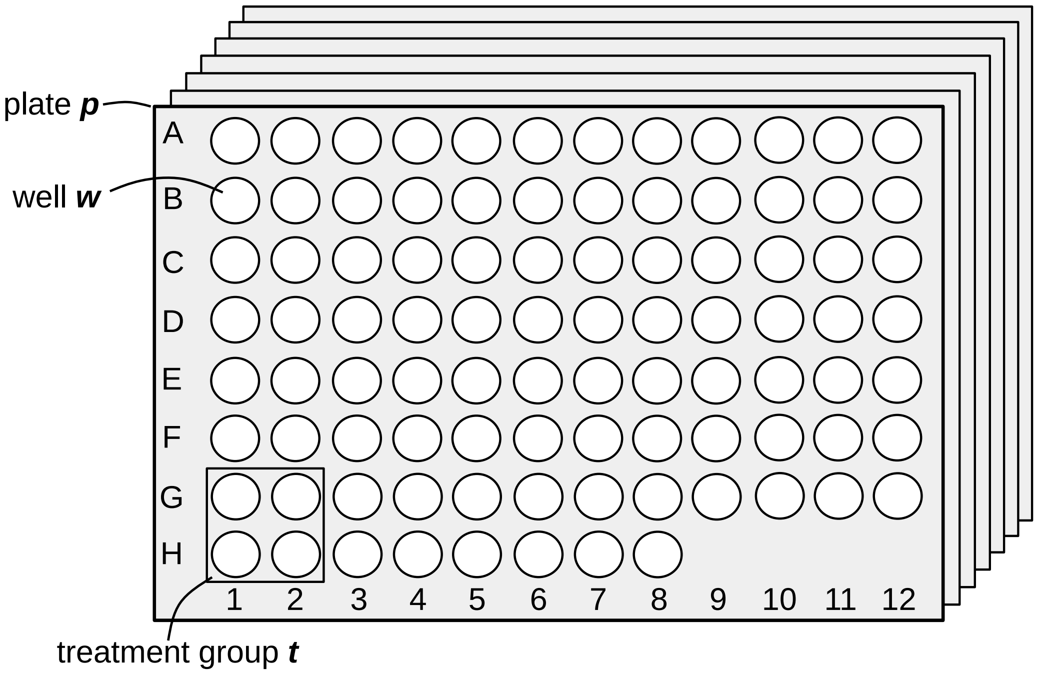


Supplementary Figure S8. Experimental design. A number of cells are seeded in well $w$, position on 96-well plate $p$. At least 4 wells on a plate are treated with treatment group $t$ (chemical compound administrated at a specific concentration).

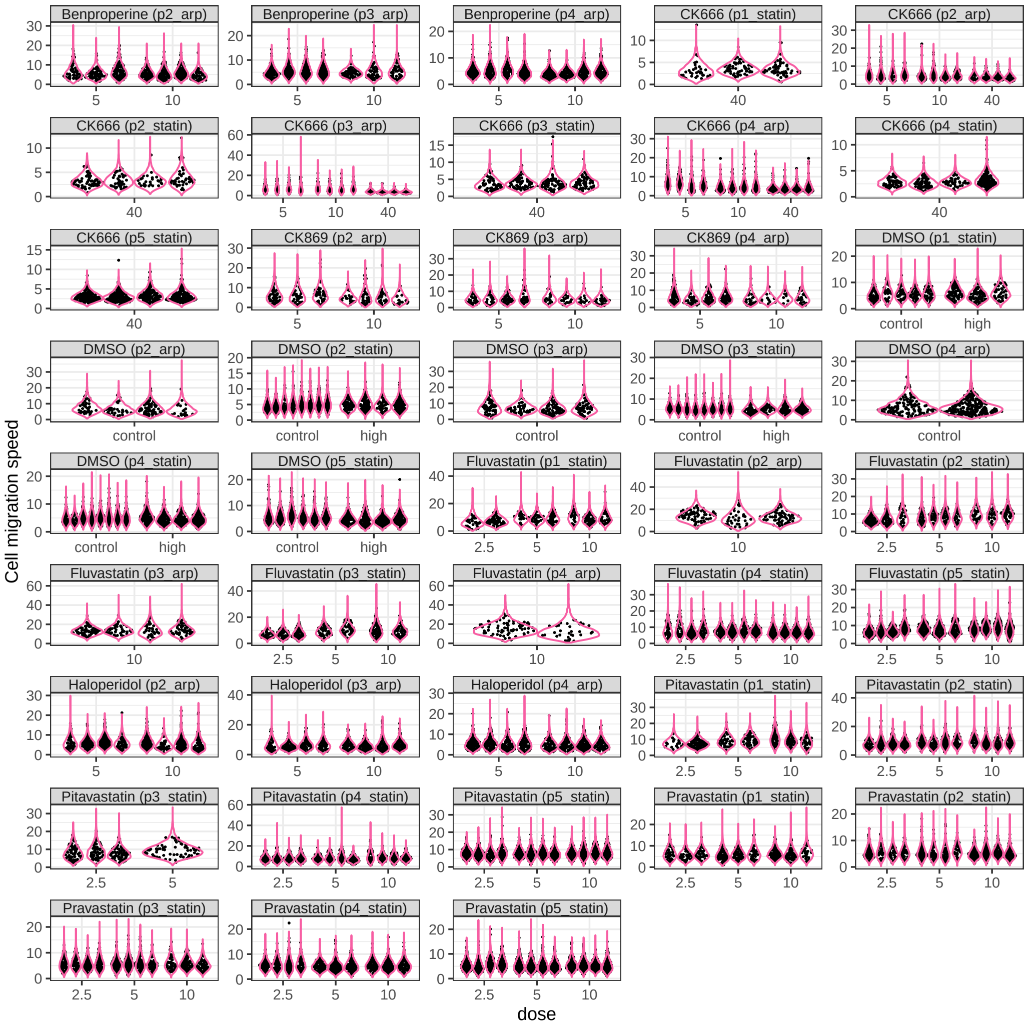


Supplementary Figure S9. Posterior predictive check of cell velocities. Each panel represents a specific chemical compound applied to wells on individual plates. Panel labels indicate the compound name and the corresponding plate identifier. Within each panel, black violins represent the distribution of observed cell velocities (y-axis), with individual cells shown as black dots. Pink violins show the model's predicted distribution of cell velocities for the same wells. The x-axis denotes the treatment dose applied to the cells in each well. The close overlap between black and pink violins indicates strong agreement between observed and predicted velocities, suggesting that the model accurately retrodicts the observed data. Panels were created using the R package ggplot2 (v3.5.2).

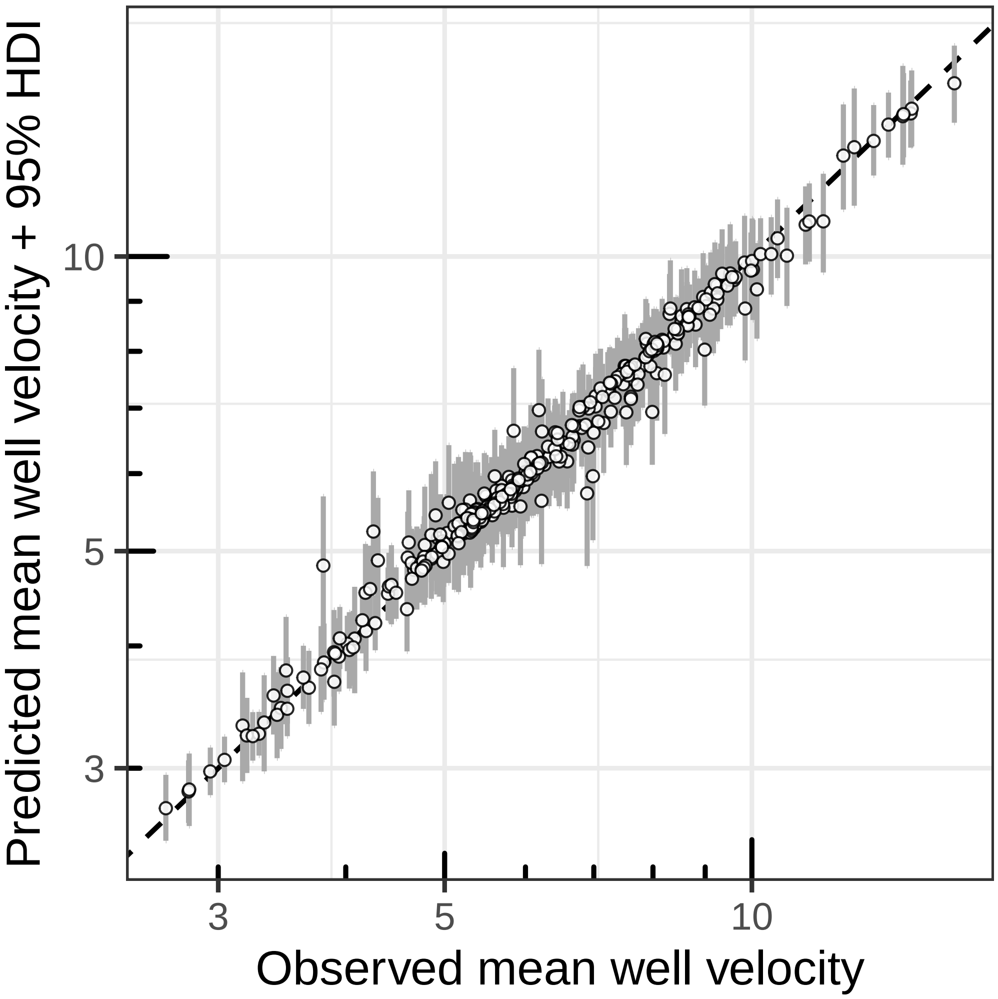


Supplementary Figure S10. Posterior predictive check of mean well velocities. Each dot represents a well, with its observed and predicted mean velocities plotted on the x- and y-axes, respectively. Gray vertical error bars indicate the 95% highest density intervals (HDIs) of the predicted mean velocities. A black dotted diagonal line marks the identity line (x = y); dots lying close to this line indicate good agreement between observed and predicted values, suggesting that the model can retrodict the observed data well. The figure was generated using the R package ggplot2 (v3.5.2).

#### References

Thomas Lin Pedersen. *patchwork: The Composer of Plots*, 2025. URL `https://patchwork.data-imaginist.com`. R package version 1.3.0.9000, https://github.com/thomasp85/patchwork.

Hadley Wickham. *ggplot2: Elegant Graphics for Data Analysis*. Springer-Verlag New York, 2016. ISBN 978-3-319-24277-4. URL `https://ggplot2.tidyverse.org`.
